# Alterations in Electroencephalography Signals in Female Fragile X Syndrome Mouse Model on a C57Bl/6J Background

**DOI:** 10.1101/2025.07.07.662836

**Authors:** Bosong Wang, Asim Ahmed, Kartikeya Murari, Ning Cheng

**Affiliations:** Faculty of Veterinary Medicine, University of Calgary, Calgary, AB, Canada; Hotchkiss Brain Institute, Cumming School of Medicine, University of Calgary, Calgary, AB, Canada; Alberta Children’s Hospital Research Institute, Cumming School of Medicine, University of Calgary, Calgary, AB, Canada; Department of Biomedical Engineering, Schulich School of Engineering, University of Calgary, Canada; Department of Electrical and Software Engineering, University of Calgary, Calgary, AB, Canada

**Keywords:** Fragile X syndrome, autism, female, mouse model, Electroencephalography, juvenile mice, C57Bl/6J, neurodevelopment, phase-amplitude and amplitude-amplitude coupling, theta-beta ratio, peak alpha frequency, signal complexity

## Abstract

**Background:** Fragile X Syndrome (FXS), the most common monogenic cause of autism spectrum disorder, arises from FMR1 gene silencing and exhibits pronounced sex differences in prevalence and phenotypic severity. Electroencephalography (EEG) has emerged as a promising translational biomarker for FXS pathophysiology, yet prior research has predominantly focused on male cohorts. In the widely used C57Bl/6J (B6) mouse strain, male *Fmr1* knockout (KO) models show increased absolute gamma power at both juvenile and adult stages, which may reflect cortical hyperexcitability. In contrast, little is known about female *Fmr1* KO mice, except that they exhibit no gamma alterations in adulthood. This gap hinders understanding of sex-specific neurodevelopmental trajectories of EEG profile in FXS. Leveraging the genetic stability and translational relevance of the B6 strain, this study compares EEG profiles between juvenile female *Fmr1* KO and wild-type (WT) B6 mice to address this critical gap.

**Methods:** Frontal-parietal differential EEG was recorded in freely behaving mice using the Open-Source Electrophysiology Recording system for Rodents. Neural activity was analyzed across three recording conditions: in the home cage, light-dark arena, and open field arena. Computed metrics included absolute/relative power, peak alpha frequency, theta-beta ratio, phase-amplitude coupling, amplitude-amplitude coupling, and multiscale entropy to assess signal complexity.

**Results:** In all recording conditions, *Fmr1* KO mice exhibited reduced absolute power in theta, alpha, and beta frequency bands compared to WT controls. Relative power analysis revealed decreased alpha activity alongside increased gamma-band power, including both low and high gamma, in the KO mice. Cross-frequency coupling was disrupted, with diminished alpha-gamma phase-amplitude coupling. Amplitude-amplitude coupling between theta or alpha and gamma power displayed distinct changes in different recording conditions. Peak alpha frequency and theta-beta ratio were both reduced or unchanged in the KO mice, depending on the recording condition. Finally, EEG signal complexity remained comparable between the two genotypes across the conditions. Behaviorally, KO mice displayed hyper-exploration in the open field test, characterized by increased center time and entries. However, no overall robust correlations between EEG power in different frequency bands and behavioral parameters in the open field test were observed.

**Discussion and Conclusion:** Our results demonstrate that juvenile female *Fmr1* KO mice on a B6 background exhibit EEG alterations highly consistent with those reported in FXS patients, particularly increased gamma and reduced alpha power. The robust increase in gamma activity reinforces its status as a reliable biomarker across preclinical and clinical studies, while alpha reductions and slowed peak alpha frequency implicate thalamocortical network involvement. Together, these findings highlight the translational value of this model for studying core circuit dysfunctions in FXS.

## 1. Introduction

Fragile X Syndrome (FXS) is the most common monogenic cause of autism spectrum disorder and intellectual disability, resulting from transcriptional silencing of the FMR1 gene and consequent loss of the Fragile X Messenger Ribonucleoprotein (FMRP) [1–3]. FXS affects approximately 1 in 4,000 males and 1 in 7,000 females, with males exhibiting more severe phenotypes possibly due to the X-linked inheritance pattern and lack of a compensatory allele [3]. Core symptoms include social interaction deficits, anxiety, hyperactivity, and cognitive impairment, with social and emotional dysfunctions being particularly prominent [4].

FMRP is a selective RNA-binding protein that critically regulates synaptic protein synthesis and neural circuit maturation [5–9]. Its absence disrupts synaptic plasticity and leads to widespread abnormalities in neural connectivity and oscillatory activity, especially in brain regions such as the prefrontal cortex, which is central to social cognition and executive function [10]. The *Fmr1* knockout (KO) mouse model recapitulates many molecular, synaptic, and behavioral features of human FXS, making it a valuable tool for mechanistic and translational research [11–14].

Neural oscillations, measured via electroencephalography (EEG), provide a non-invasive window into observing large-scale brain network dynamics and have emerged as promising translational biomarkers for neurodevelopmental disorders, including FXS [15, 16]. EEG captures synchronized activity across frequency bands—delta, theta, alpha, beta, and gamma—each associated with distinct cognitive and physiological functions [16–20]. Previous studies in FXS have consistently reported alterations in EEG power spectra, particularly increased gamma power and disrupted low-frequency rhythms, which are thought to reflect underlying excitatory-inhibitory (E/I) imbalance and thalamocortical dysregulation [21–23].

Despite significant progress, important gaps remain. Most EEG studies in FXS have focused on male subjects, both in clinical and preclinical settings, revealing robust increases in absolute and relative gamma power in male *Fmr1* KO mice across both juvenile and adult ages [24–28]. In contrast, female *Fmr1* KO mice on the C57Bl/6J (B6) background show no gamma power changes in adulthood (findings in other frequency bands were not reported), and the juvenile EEG phenotype remains uncharacterized [27]. Given the pronounced sex differences in FXS at molecular, circuit, and behavioral levels, and the widespread use of the B6 strain in neuroscience research, there is a critical need to investigate neural oscillatory abnormalities in juvenile female models [29–31].

Furthermore, advanced EEG analytical approaches—including assessments of peak alpha frequency (PAF), theta-beta ratio (TBR), cross-frequency coupling, and signal complexity—offer deeper insights into network dysfunctions beyond traditional power spectral analysis. These metrics have shown promise for identifying circuit-level biomarkers relevant to cognitive and behavioral changes in FXS and autism, and related neurodevelopmental disorders [32–35].

The present study addresses these gaps by systematically characterizing EEG oscillatory activity and its behavioral correlations in juvenile female *Fmr1* KO mice on the B6 background. By employing a comprehensive suite of EEG analyses across multiple behavioral paradigms, this work aims to help elucidate sex-specific neural signatures of FXS, inform biomarker development, and advance understanding of the neurophysiological mechanisms underlying social and cognitive changes in this disorder.

## 2. Materials and Methods

### 2.1 Animals

The *Fmr1* KO mice used in this study were generated by mating homozygous *Fmr1* knockout females (*Fmr1 -/-*) with hemizygous *Fmr1* knockout males (*Fmr1 -/y*). Breeder wild-type (WT) mice (C57BL/6J, Jackson Laboratory stock No: 000664) and *Fmr1* knockout mice (B6.129P2-*Fmr1*^tm1Cgr^/J, Jackson Laboratory stock No: 003025) were sourced from the Jackson Laboratory (ME, USA). Mice were housed at the animal facility of the Cumming School of Medicine, University of Calgary. They were group-housed with up to five same-sex littermates per cage. The housing environment has a controlled 12-hour light-dark cycle (lights on at 07:00 AM) with ad libitum access to food and water. Pups were weaned at approximately 21 days of age and fed on a standard mouse chow diet. Testing was conducted on female *Fmr1* homozygous knockout and wild-type mice bred in-house at the University of Calgary. Testing was performed between 09:00 and 19:00 hours.

### 2.2 Ethics

All procedures in this study were performed in accordance with the recommendations of the Canadian Council for Animal Care. The protocol of this study was approved by the Health Sciences Animal Care Committee of the University of Calgary.

### 2.3 EEG surgery

EEG surgery was performed on mice using established protocols [36–38]. Anesthesia was induced with 5% isoflurane and maintained at 1%–2% isoflurane throughout the procedure. Mice were positioned on a heated pad attached to stereotaxic equipment to maintain body temperature. To prevent desiccation, the eyes were protected with eye ointment. Following shaving, the scalp was cleaned with 70% ethanol and iodine, and lidocaine was applied to minimize local pain. Once the absence of a toe pinch reflex confirmed adequate anesthesia, the top layer of skin was removed, and the muscles and connective tissue were carefully retracted to expose the skull. The skull was further cleaned with hydrogen peroxide to ensure removal of any remaining connective tissue. For differential frontal-parietal recordings, three small burr holes were drilled into the skull at the following coordinates relative to bregma: ground (AP: +2.5 mm, ML: +1.3 mm), left parietal cortex (AP: −2.2 mm, ML: −2.5 mm), and left frontal cortex (AP: +2.5 mm, ML: −1.3 mm). The electrodes were then inserted into these holes and connected to a miniature connector. Dental cement was used to secure the electrode assembly to the skull. Following EEG surgery, mice received analgesic medication and were maintained on a heating pad until full ambulation recovery.

### 2.4 EEG signal recording and analysis

All behavioral testing and EEG recordings were conducted 5-7 days post-surgery to ensure adequate recovery. For EEG recordings, mice underwent brief isoflurane anesthesia for connection to a stand-alone EEG system. Each recording session comprised a 30-minute baseline in the home cage followed by a 30-minute recording in the behavioral test chambers. This protocol was repeated across two consecutive days, each using a different behavioral paradigm: Light-Dark (LD) test and Open Field Test (OFT).

EEG signals exhibiting abnormally large amplitudes or saturation artifacts, likely resulting from post-surgical electrode-connector interface issues, were excluded from analysis.

EEG data analysis followed established methodologies [37, 38]. After verifying recording integrity through visual inspection, 5-minute artifact-free segments were selected near the end of each experimental condition (home cage, Light-Dark and Open Field). These segments were selected from periods of confirmed wakefulness, as determined by simultaneous behavioral monitoring. Spectral decomposition was performed using MATLAB R2023a (MathWorks). Signals were segmented into 2-second epochs with 50% overlap and processed using Welch’s modified periodogram method, yielding a spectral resolution of 0.5 Hz. Power spectral density (PSD) estimates were computed by averaging periodograms across epochs. Band-specific power was quantified through trapezoidal numerical integration of the PSD across each frequency band: delta (1–4 Hz), theta (4–8 Hz), alpha (8–13 Hz), beta (13–30 Hz), and gamma (30–100 Hz). The gamma band was further subdivided into low-gamma (30–60 Hz) and high-gamma (60–100 Hz) components. Relative band power was calculated as the ratio of the power within a specific frequency band to the sum of power across all frequency bands (delta, theta, alpha, beta, and gamma).

### 2.5 Behavioral tests

The LD test leverages rodents’ natural aversion to bright light and their exploratory behavior in response to a novel environment [39]. The testing arena consists of a dark chamber and an adjacent brightly lit chamber. During the experiment, mice were allowed to freely explore this two-compartment arena for 10 minutes. The number of entries into each compartment, duration spent in each compartment, and latency to enter the light compartment were quantified.

The OFT is a widely used platform for assessing locomotor activity and anxiety-like behaviors in animal models [40]. The experiment involves placing subjects in a novel square arena under standard room illumination for 10 minutes, during which locomotor activity and center zone exploration were quantified.

All testing sessions were recorded using Ethovision (Noldus). Between each test, the chamber was thoroughly cleaned with 70% ethanol and allowed to dry for several minutes to ensure standardized conditions for subsequent tests.

### 2.6 PAF

To determine the PAF for each mouse, we analyzed the PSD values within the alpha frequency range (8–13 Hz). PSD was estimated with a spectral resolution of 0.1 Hz, resulting in 51 discrete frequency bins across the specified range. The frequency corresponding to the highest PSD value within this range was identified as the PAF. This calculation was performed individually for each mouse in both WT and KO groups.

### 2.7 TBR

TBR was determined by dividing the absolute theta power by the absolute beta power for each animal.

### 2.8 Phase – Amplitude Coupling (PAC)

To investigate cross-frequency interactions, we analyzed PAC between gamma oscillations (30– 100 Hz) and lower-frequency rhythms within the theta (4–8 Hz) and alpha (8–13 Hz) bands. For this analysis, 2-minute epochs of artifact-free spontaneous EEG activity were processed with the reduced interference distribution-Rihaczek time-frequency distribution, which simultaneously extracts phase time series from theta/alpha oscillations and amplitude envelopes from gamma activity [41]. Coupling strength was quantified using the mean vector length (MVL) modulation index, calculated as the circular consistency between normalized gamma amplitude and instantaneous theta/alpha phase [42]. Phase frequencies were assessed at 1 Hz intervals across the theta (4–8 Hz) and alpha (8–13 Hz) bands, paired with amplitude frequencies spanning the gamma range (30–100 Hz) in 2 Hz increments. Normalized PAC strength was derived using a block-swapping surrogate data approach.

### 2.9 Amplitude – Amplitude Coupling (AAC)

To investigate interdependencies between oscillatory power across frequency bands, we conducted cross-frequency AAC analyses [24]. These examined relationships between theta (4–8 Hz), lower alpha (8–10 Hz), and upper alpha (10–13 Hz) oscillations with both low gamma (30–60 Hz) and high gamma (60–100 Hz) activity. EEG time series were segmented into consecutive 2-second epochs for analysis. For each subject, Spearman’s rank correlation coefficients were computed for all pairwise combinations of low-frequency (theta, lower alpha, upper alpha) and high-frequency (low gamma, high gamma) bands [24]. This non-parametric approach accounted for potential non-normal distributions in power time courses. Correlation coefficients were subsequently converted to normally distributed z-scores using Fisher’s Z-transformation to enable group-level comparisons.

### 2.10 Multi-Scale Entropy (MSE)

MSE was employed to quantify signal complexity in EEG recordings. To compute MSE in mice, we followed the established method [43]. The analysis consisted of two primary stages. First, the original EEG signal was segmented into 2-second epochs and subjected to a coarse-graining procedure to generate scaled time series. For each scale factor (τ = 1–40), non-overlapping windows of length τ were averaged, progressively shortening the time series as τ increased. Second, sample entropy (*SampEn*) was computed for each scaled time series to assess signal irregularity [44]. *SampEn* parameters were set to an embedding dimension (m = 2) and tolerance threshold (r = 0.5), where r represented 50% of the epoch’s standard deviation. This metric calculates the negative natural logarithm of the conditional probability that sequences matching for *m* points remain similar at *m+1* points, where A and B denote matches for m+1 and m points, respectively. MSE values were averaged across all epochs within the 5-minute recording for each mouse, yielding scale-specific complexity measures (τ = 1–40).

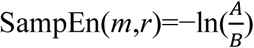

To evaluate overall signal complexity, the area under the MSE curve (AUC) across scales 1–40 was computed as the Complexity Index. For granular analysis, scales were divided into short-term (1–20) and long-term (21–40) dynamics, with group averages calculated for each range.

### 2.11 Statistical tests and analysis

Statistical analyses were conducted using GraphPad Prism (version 10, GraphPad Software, San Diego, CA, USA). All data are presented as mean ± standard error of the mean (SEM). The threshold for statistical significance was set at α = 0.05 for all analyses. Between-group comparisons were performed using unpaired two-tailed t-tests, with genotype as the primary independent variable. Normality of data distribution was assessed using the Shapiro-Wilk test. In cases where parametric assumptions were violated, Mann-Whitney U tests were applied.

For correlation analyses, Spearman’s rank correlation coefficients were calculated for AAC measures, while Pearson’s correlation coefficients were used to assess linear relationships between EEG parameters and behavioral outcomes. To control for multiple comparisons in the correlation analyses, the p values were corrected with false discovery rate using the Benjamini-Hochberg procedure.

## 3. Results

### 3.1 Reductions in Lower-Frequency Absolute EEG Power in *Fmr1* KO Mice

We observed genotype-dependent differences in absolute spectral power across behavioral paradigms. During the home cage recordings, *Fmr1* KO mice exhibited significant reductions in absolute power in delta, theta, alpha, and beta frequency bands compared to WT controls **(Fig. 1)**. No genotype-driven differences were observed in the gamma band. Total EEG power was also markedly lower in KO mice.

**Figure 1.**
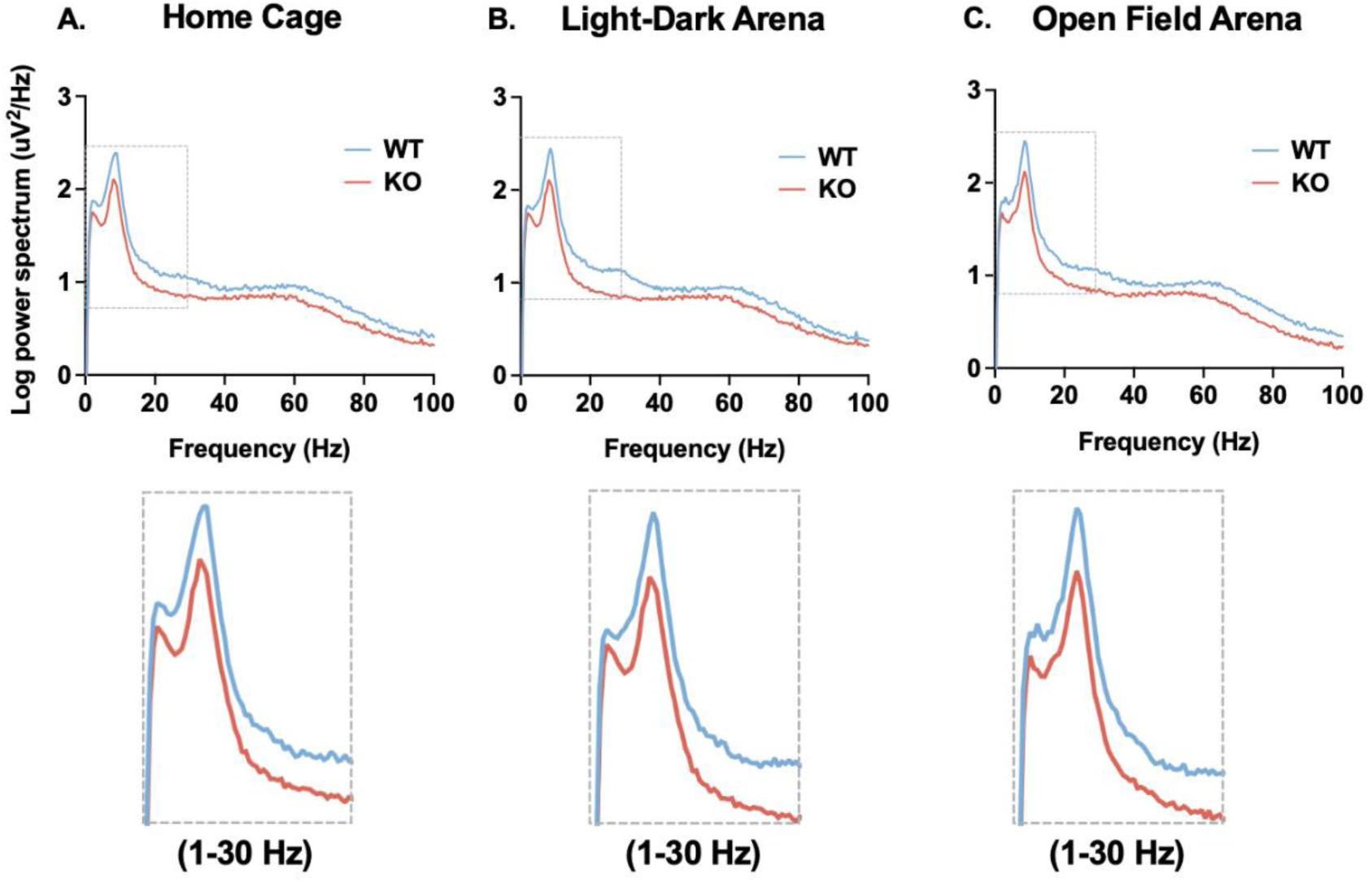
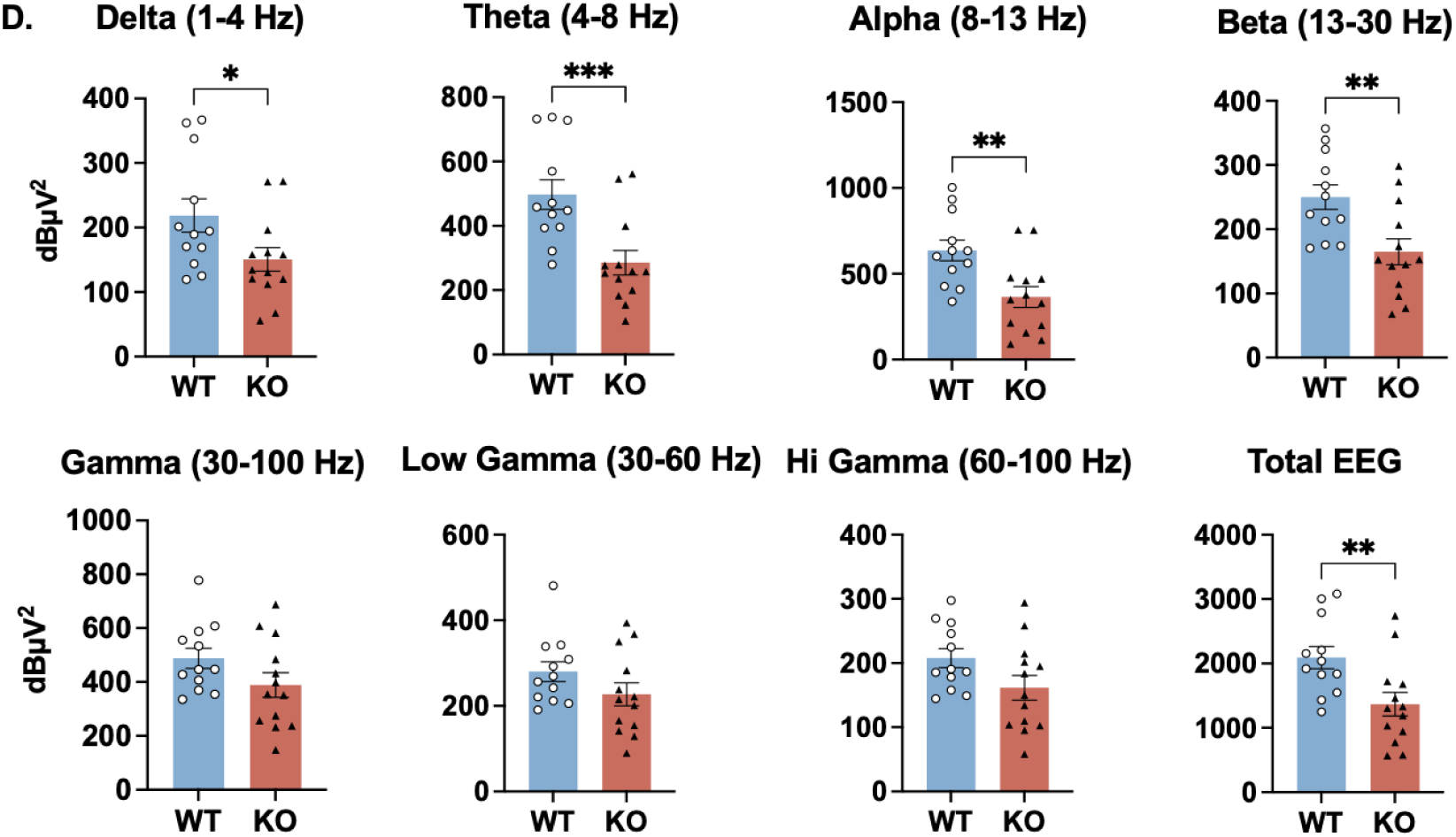
Absolute EEG power in KO compared to WT mice. (A, B, & C) Power spectra of KO and WT mice in the home cage, LD, and OFT. (D) Comparison of absolute EEG power between WT and KO, in delta, theta, alpha, beta, and gamma, low and high gamma frequency bands, and total power in home cage recordings. Data are presented as mean ± SEM for each group (WT: blue; KO: red). Two-tailed unpaired t-tests were performed between WT and KO groups, with significance indicated as following: *: p < 0.05, **: p < 0.01, ***: p < 0.001.

During the LD and OFT, similar patterns emerged: KO mice displayed decreased absolute power in theta, alpha, beta, and high gamma bands, alongside reduced total power **(Table I)**. In contrast to home cage findings, delta power remained comparable between genotypes in both LD and OFT, while significant reduction in high gamma sub-band activity was observed.

**Table I.** P-value summary of comparing absolute EEG power in light-dark and open field arenas between female juvenile WT and KO mice.

| Frequency bands | Home Cage | Light-Dark Arena | Open Field Arena |
| --- | --- | --- | --- |
| Delta | **0.0398** ↓ | 0.2254 | 0.3567 |
| Theta | **0.0010** ↓ | **0.0030** ↓ | **0.0083** ↓ |
| Alpha | **0.0044** ↓ | **0.0086** ↓ | **0.0046** ↓ |
| Beta | **0.0055** ↓ | **0.0036** ↓ | **0.0089** ↓ |
| Gamma | 0.1098 | 0.0747 | 0.0909 |
| Low Gamma | 0.1499 | 0.1235 | 0.1544 |
| High Gamma | 0.0744 | **0.0356** ↓ | **0.0380** ↓ |
| Total EEG power | **0.0089** ↓ | **0.0123** ↓ | **0.0100** ↓ |

↓ indicates that EEG power in KO mice is significantly lower than in WT mice for the corresponding frequency band and condition

### 3.2 Altered Relative Power in Theta, Alpha, and Gamma Bands in *Fmr1* KO Mice

Relative spectral power analysis revealed genotype-specific shifts in oscillatory profiles. During home cage and LD recordings, *Fmr1* KO exhibited significant reductions in theta and alpha power alongside increased gamma activity, including both low and high gamma, compared to WT controls **(Fig. 2)**. In the OFT, similar differences were observed, except that theta power differences were absent **(Table II)**. No genotype differences emerged in delta or beta bands in any conditions.

**Figure 2.**
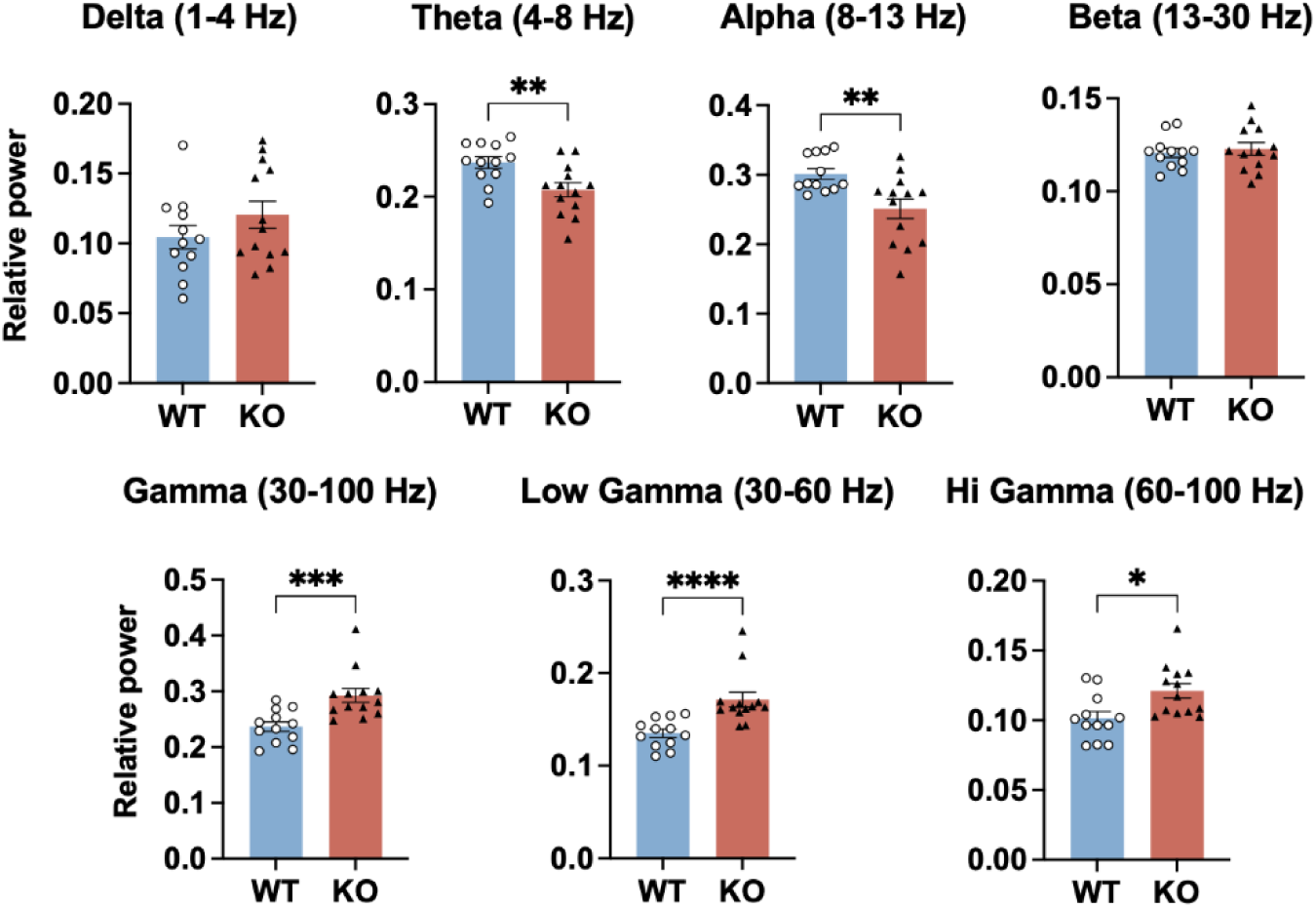
Relative theta and alpha power decreased with relative gamma power increased in KO mice. Figure shows relative EEG power for delta, theta, alpha, beta, and gamma, including low and high gamma, in home cage recording. Data are presented as mean ± SEM for each group (WT: blue; KO: red). Two-tailed unpaired t-tests were performed between WT and KO groups, with significance indicated as following: *: p < 0.05, **: p < 0.01, ***: p < 0.001, ****: p < 0.0001.

**Table II.**
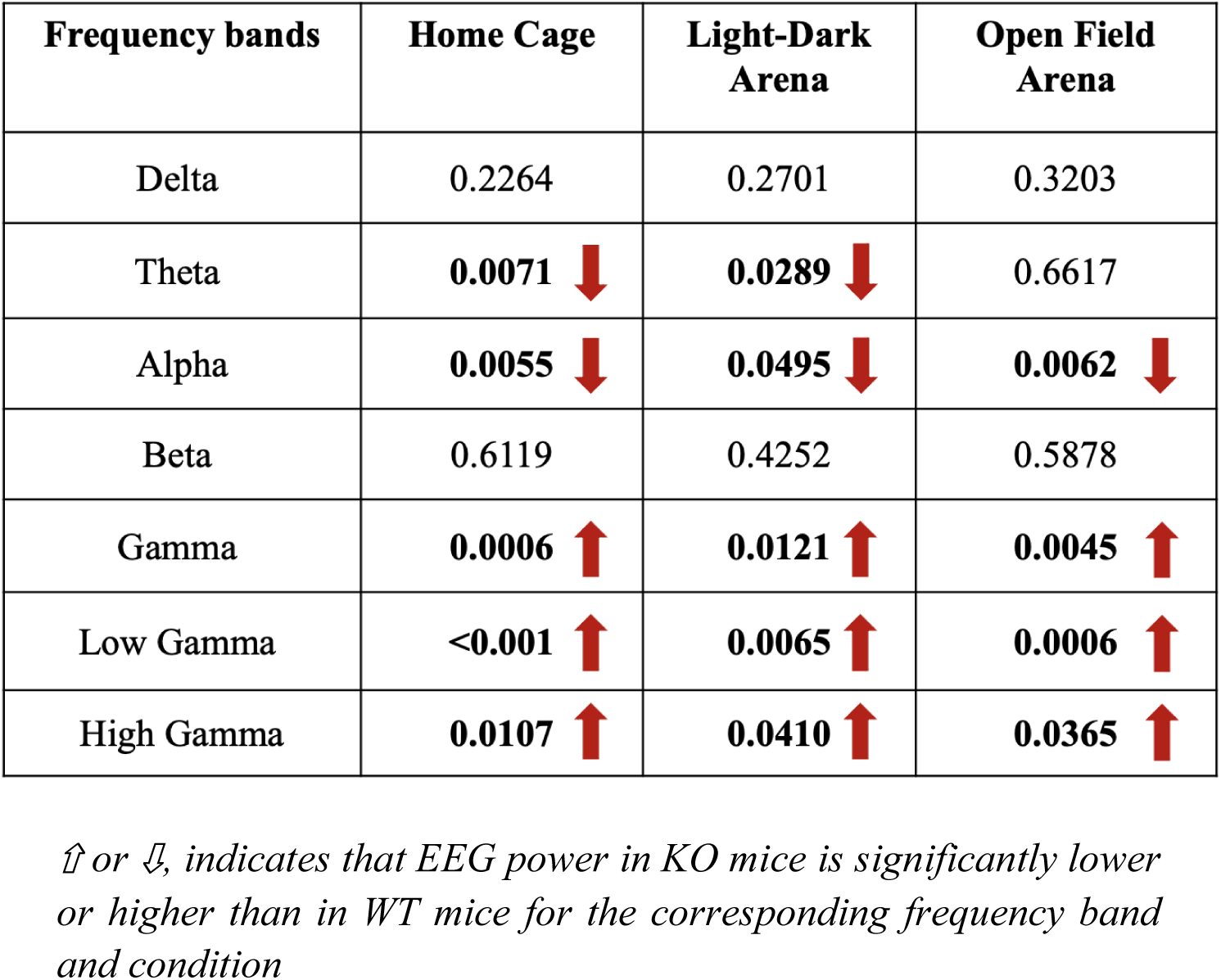
P-value summary of comparing relative EEG power in light-dark and open field arenas between female juvenile Fmr1 KO and WT mice.

### 3.3 Home Cage Specific Reduction in PAF

PAF is defined as the dominant oscillatory rhythm within the alpha band, reflects thalamocortical network integrity, and is implicated in cognitive processing [32]. Its dynamic is linked to cognitive processing and neurodevelopmental maturation [45]. In this study, PAF was quantified during home cage, LD, and OFT paradigms. Our results revealed that *Fmr1* KO mice exhibited a significant reduction in PAF during home cage recordings compared to WT controls **(Fig. 3A)**. Notably, this reduction was state-specific: no genotype differences emerged in PAF during LD or OFT.

**Figure 3.**
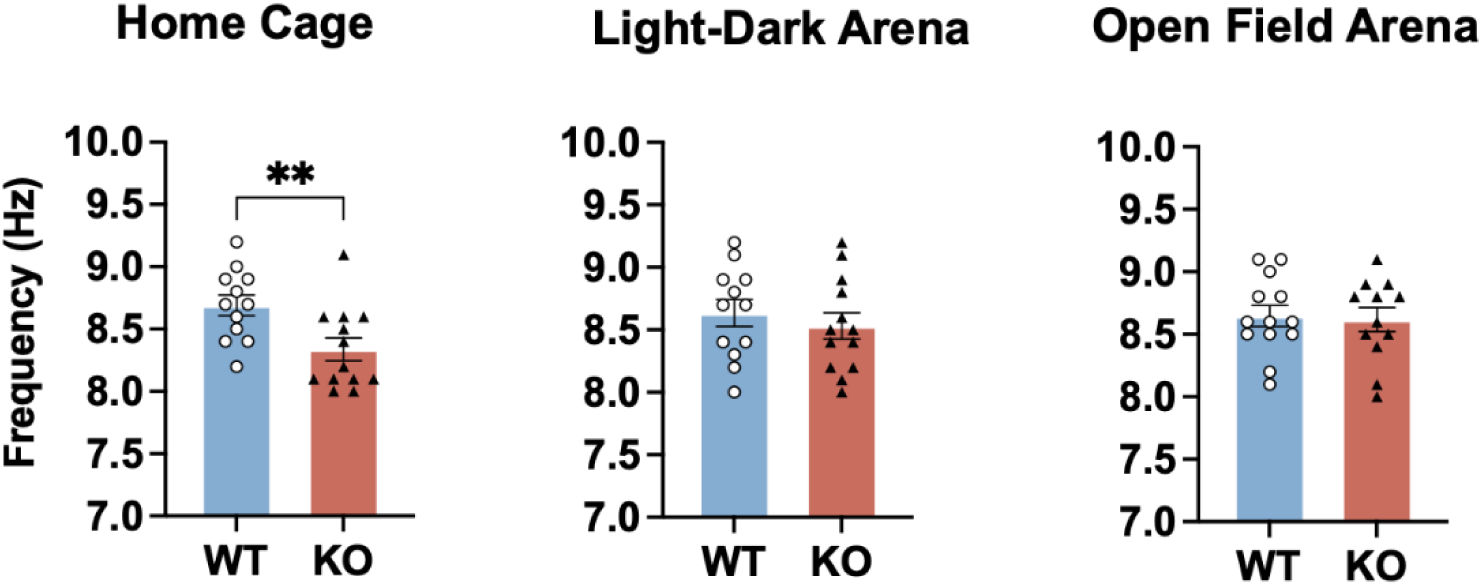
PAF decreased in KO mice in home cage recordings. Comparison of PAF between WT and KO mice in the home cage, LD and OFT. Data are presented as mean ± SEM for each group (WT: blue; KO: red). Two-tailed unpaired t-tests were performed between WT and KO groups, with significance indicated as following: **: *p* < 0.01.

### 3.4 Decreased TBR During Home Cage and Light-Dark Test in *Fmr1* KO Mice

TBR was calculated as the ratio of theta to beta power. It is well-established for attentional processes and neurodevelopmental disorders such as ADHD [33, 46]. Elevated TBR may reflect cortical hypo-arousal and is linked to reduced cognition and inattention [33]. In the present study, TBR was significantly reduced in *Fmr1* KO mice during home cage recordings and LD compared to WT controls **(Fig. 4)**. This reduction stemmed from genotype-specific decreases in relative theta power (p < 0.05) alongside stable beta power (p > 0.1). No TBR differences emerged in the OFT.

**Figure 4.**
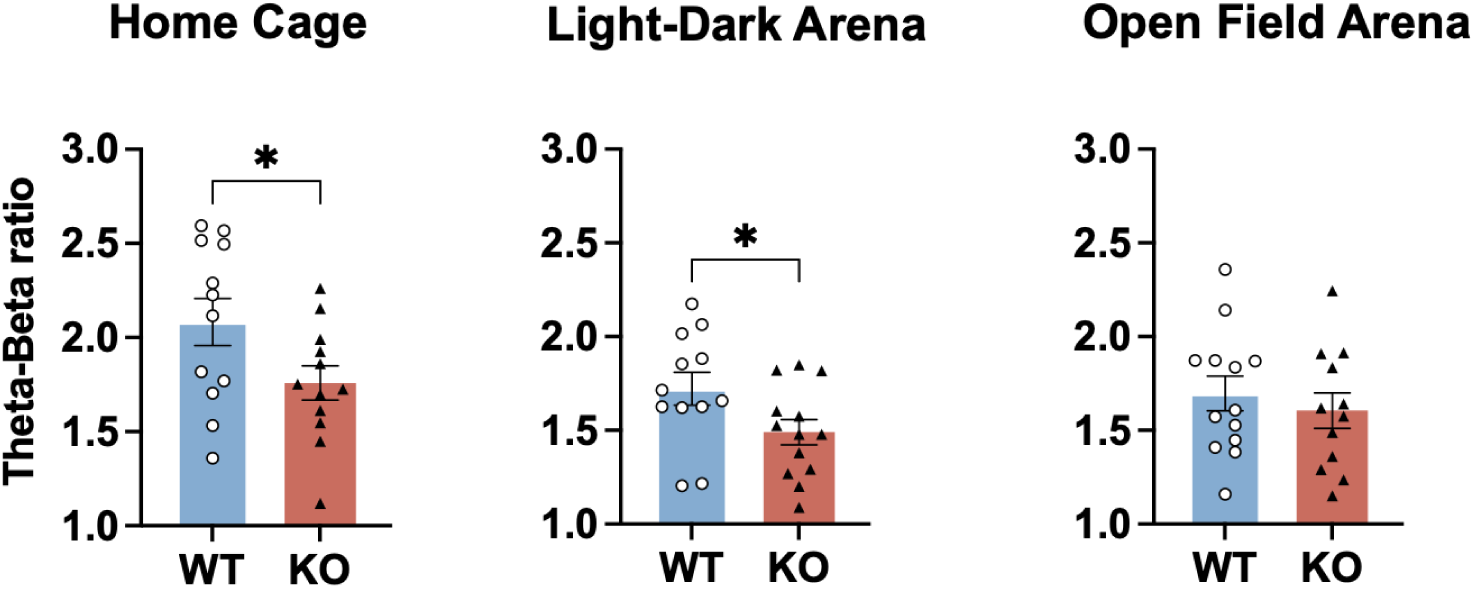
TBR decreased in KO compared to WT mice in home cage and LD. Comparison of TBR between WT and KO mice in the home cage, light-dark and open field arenas. Data are presented as mean ± SEM for each group (WT: blue; KO: red). Two-tailed unpaired t-tests were performed between WT and KO groups, with significance indicated as following: *: *p* < 0.05.

### 3.5 Disrupted Cross Frequency PAC in *Fmr1* KO Mice Across Experimental Conditions

The phase of low-frequency oscillations modulates the amplitude of high-frequency activity and serves as a mechanism for integrating spatiotemporal neural processes [34]. In sensory systems, PAC coordinates local computations (e.g., gamma oscillations) with broader network dynamics (e.g., theta/alpha rhythms), facilitating perceptual binding and top-down prediction-processes often impaired in neurodevelopmental disorders [34]. Theta-gamma PAC is critical for organizing working memory and encoding long-term memory, while alpha-gamma PAC facilitates attentional focus and sensory integration [47–49]. Given the central role of these processes in FXS-related cognitive deficits, we analyzed theta- and alpha-gamma PAC separately across behavioral states. We found *Fmr1* KO mice exhibited PAC disruptions. Specifically, both alpha-gamma and theta-gamma PAC were reduced in the KO mice compared to WT controls, except during home cage recordings, only alpha-gamma, but not theta-gamma PAC was reduced **(Fig. 5)**.

**Figure 5.**
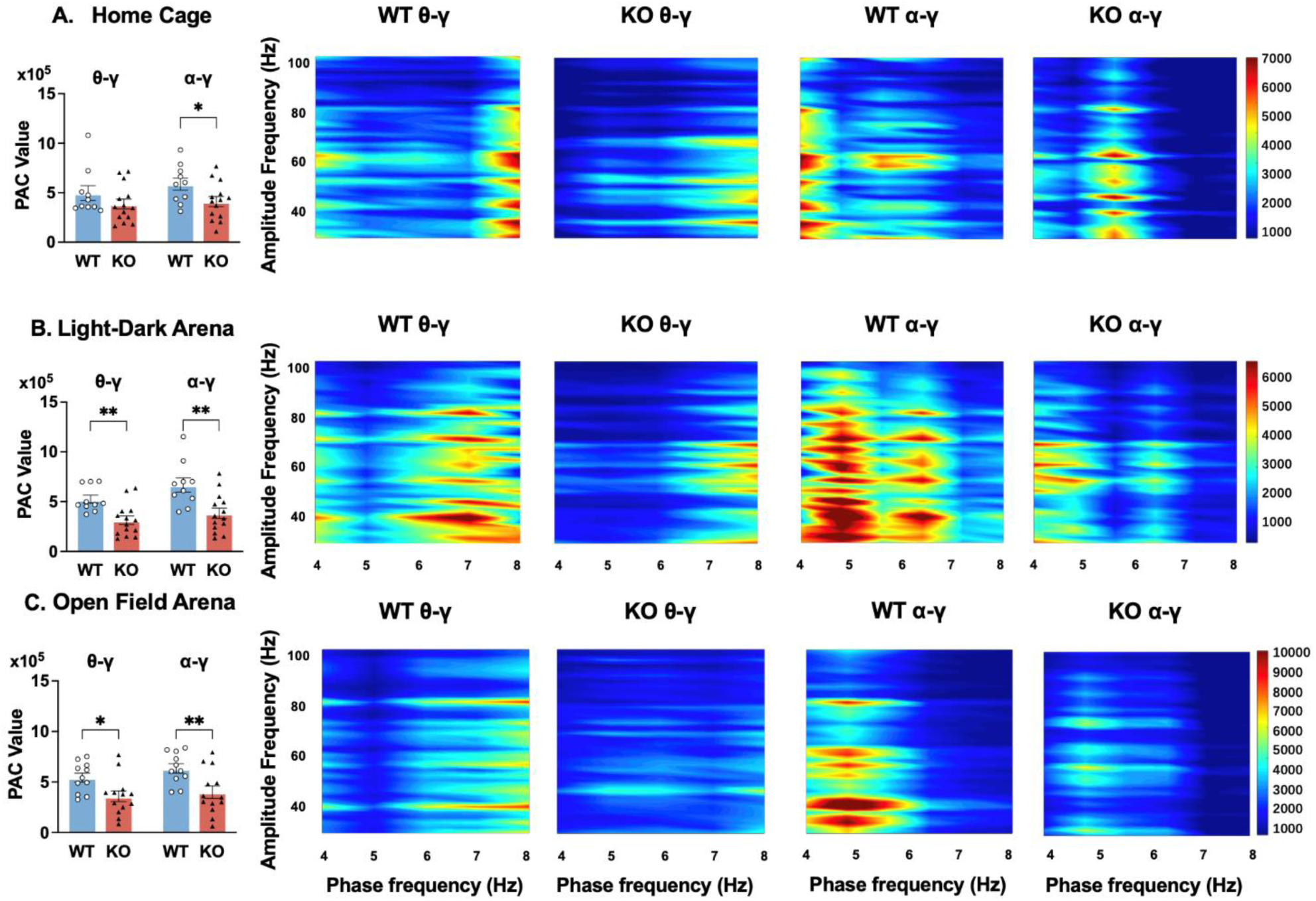
PAC was generally decreased in KO mice compared to WT controls across conditions, for both theta-gamma coupling and alpha-gamma coupling. (A, B, & C) Shows the quantification of PAC and phase-amplitude map for recordings in home cage, light-dark, and open field arenas. The left panels show the quantification of total MVL values and comparison between WT and KO mice. The right panels show representative phase-amplitude heat map between the amplitude of the gamma band (30 – 100 Hz, amplitude-frequency) and the phase of the low-frequency band (theta 4 – 8 Hz, alpha 8-13 Hz, phase-frequency). The color scale is representative of MVL values, a measurement of coupling strength. Data are presented as mean ± SEM for each group (WT: blue; KO: red). Two-tailed unpaired t-tests were performed between WT and KO groups, with significance indicated as follows: *: *p* < 0.05, **: *p* < 0.01.

### 3.6 Alterations in Cross Frequency AAC Patterns in *Fmr1* KO Mice

Another method to evaluate the potential association of different frequency bands, particularly their amplitude, is AAC. It quantifies the temporal co-modulation of power between distinct frequency bands. AAC captures coordinated intensity fluctuations critical for large-scale network communication [35]. For the current study, we analyzed six AAC pairs based on their established roles in cognitive processes and prior findings in FXS models [22]. During home cage recordings, *Fmr1* KO mice exhibited increased theta–high gamma and reduced alpha–high gamma coupling compared to WT controls **(Fig. 6)**. No genotype differences emerged in LD. During OFT, KO mice displayed heightened theta–low gamma AAC (**Tabel III).**

**Figure 6.**
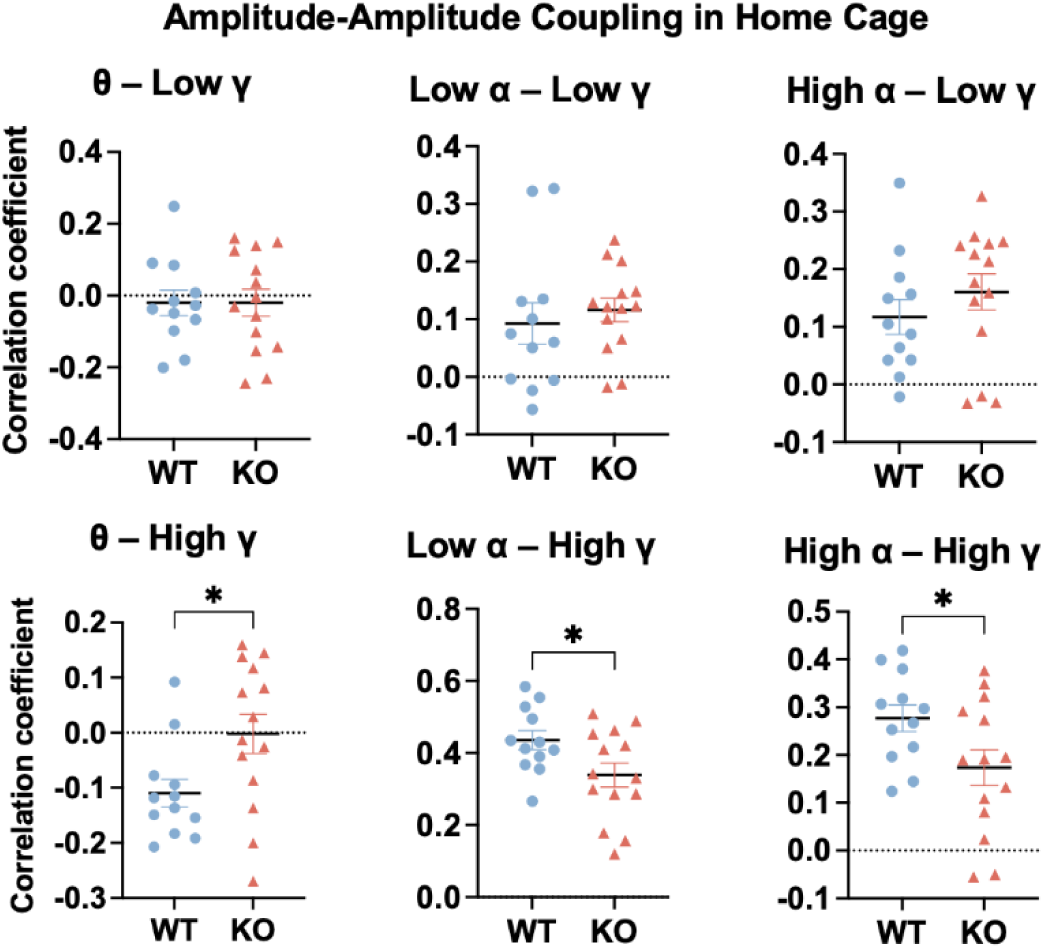
AAC between low frequency (theta and alpha) and high frequency power (gamma) in home cage. Cross frequency AAC including theta-low gamma, theta-high gamma, low alpha-low gamma, low alpha-high gamma, high alpha-low gamma, and high alpha-high gamma are compared between WT and KO mice in home cage, light-dark and open field arenas. Data are presented as mean ± SEM for each group (WT: blue; KO: red). Two-tailed unpaired t-tests were performed between WT and KO groups, with significance indicated as following: *: *p* < 0.05.

**Table III.** P-value summary of comparing six different AAC across behavioral paradigms between female juvenile KO and WT mice.

| Analysis | Condition | p value<br>(WT vs KO) | Analysis | Condition | p value<br>(WT vs KO) | Analysis | Condition | p value<br>(WT vs KO) |
| --- | --- | --- | --- | --- | --- | --- | --- | --- |
| **Home**<br>**Cage** | $\theta$ – Low $\gamma$ | 0.9933 | **Light-**<br>**Dark**<br>**Arena** | $\theta$ – Low $\gamma$ | 0.8173 | **Open**<br>**Field**<br>**Arena** | $\theta$ – Low $\gamma$ | **0.0223** ↑ |
| | $\theta$ – High $\gamma$ | **0.0253** ↑ | | $\theta$ – High $\gamma$ | 0.4777 | | $\theta$ – High $\gamma$ | 0.1008 |
| | Low $\alpha$ – Low $\gamma$ | 0.5597 | | Low $\alpha$ – Low $\gamma$ | 0.8374 | | Low $\alpha$ – Low $\gamma$ | 0.5653 |
| | Low $\alpha$ – High $\gamma$ | **0.0370** ↓ | | Low $\alpha$ – High $\gamma$ | 0.3160 | | Low $\alpha$ – High $\gamma$ | 0.3195 |
| | High $\alpha$ – Low $\gamma$ | 0.3293 | | High $\alpha$ – Low $\gamma$ | 0.5615 | | High $\alpha$ – Low $\gamma$ | 0.8338 |
| | High $\alpha$ – High $\gamma$ | **0.0423** ↓ | | High $\alpha$ – High $\gamma$ | 0.8456 | | High $\alpha$ – High $\gamma$ | 0.3598 |

↑ or ↓, indicates that EEG power in KO mice is significantly lower or higher than in WT mice for the corresponding frequency band and condition

### 3.7 MSE Analysis Reveals No Significant Difference Between KO and WT Mice

Signal complexity, quantified via MSE, reflects the temporal richness of neural activity and is implicated in brain maturation, cognitive function, and neurodevelopmental disorders [43]. MSE measures entropy across multiple time scales, where lower scales (1–20) reflect local neural dynamics and higher scales (21–40) capture long-range network interactions.

In this study, MSE analysis revealed no genotype differences in *Fmr1* KO versus WT mice. Complexity indices (area under MSE curves for scales 1–40) were comparable between KO and WT mice during recordings in home cage, LD, and OFT **(Fig. 7A)**. Subscale analysis of shorter (1–20) and longer (21–40) time scale also showed no significant differences **(Fig. 7B–C)**.

**Figure 7.**
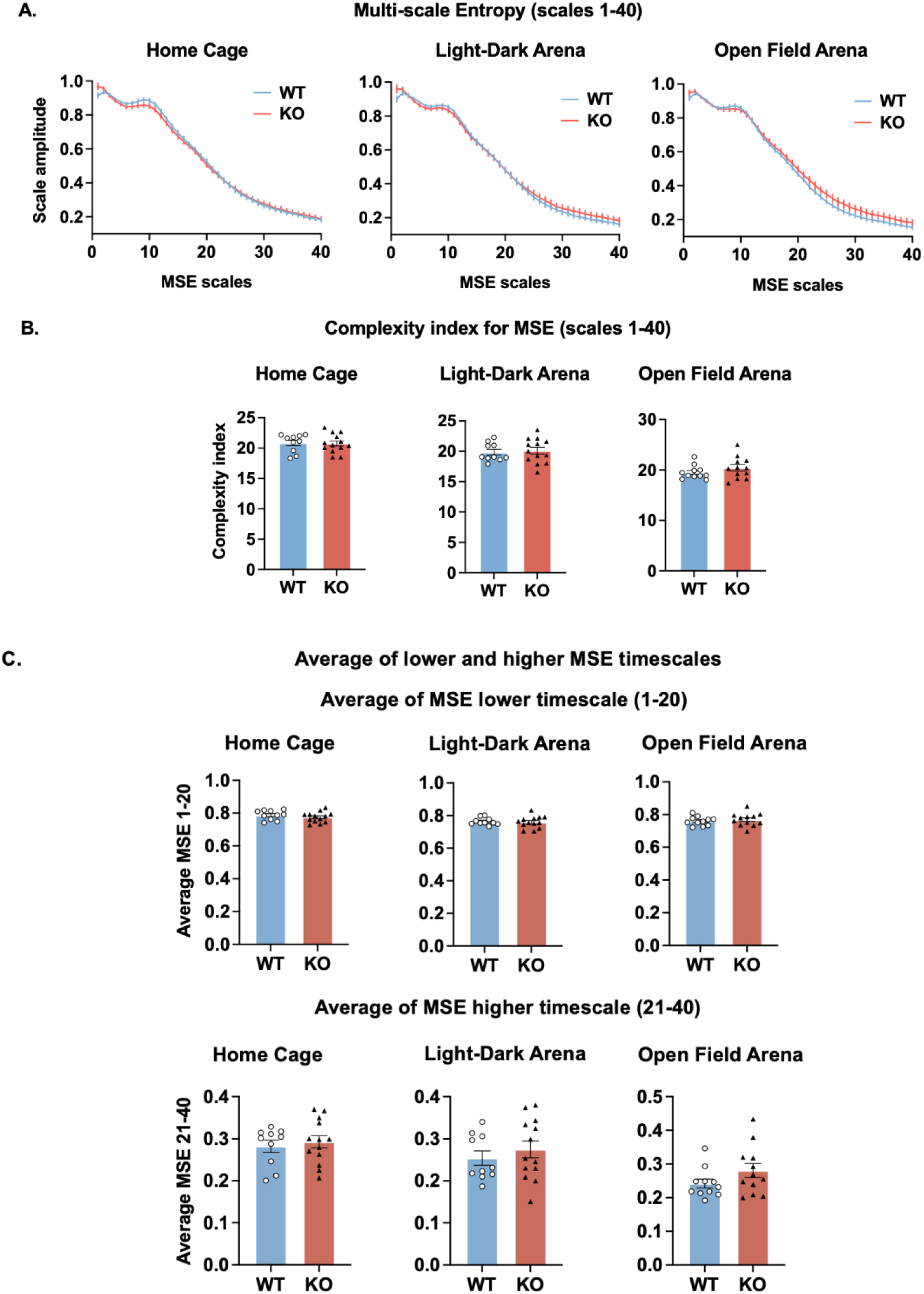
MSE analysis in WT and KO mice. (A) Average amplitude of MSE at timescales (1-40) for WT and KO mice. (B) Complexity index (area under the curve) for MSE (1-40) was compared between WT and KO mice in home cage, LD and OFT. (C) Average of MSE lower timescale (MSE 1-20) and higher timescale (MSE 21-40), compared between WT and KO mice in home cage, LD and OFT. Data are presented as mean ± SEM for each group (WT: blue; KO: red). Two-tailed unpaired t-tests were performed between WT and KO groups.

### 3.8 Enhanced Center-Directed Exploration in *Fmr1* KO Mice During Open Field Test

Behavioral analysis revealed genotype-specific differences in exploratory patterns during the OFT. *Fmr1* KO mice exhibited significantly increased center zone exploration compared to WT controls, as evidenced by three key metrics: greater distance traveled in the center zone, higher frequency of center entries, and prolonged cumulative time spent in the center zone **(Fig. 8)**. In contrast, no behavioral differences emerged in the LD, where both genotypes displayed comparable latencies to enter the light compartment, time spent in the light zone, distance travelled in the light compartment and velocity in the light compartment **(Supplementary Fig.1)**.

**Figure 8.**
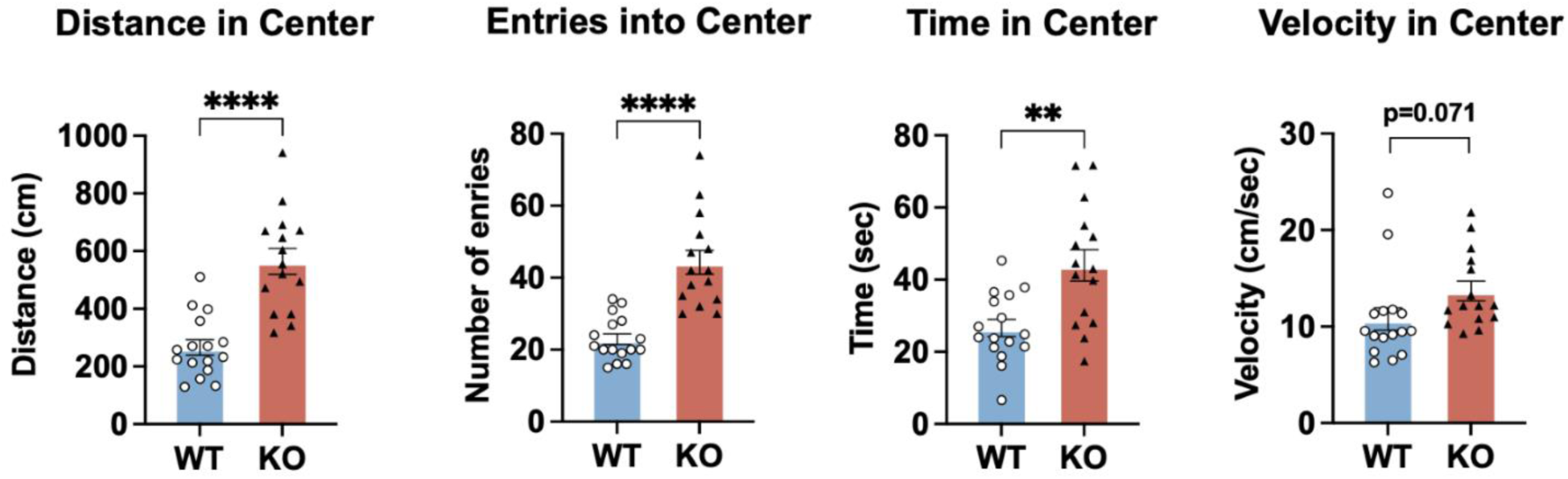
OFT parameters measured in the center zone for WT and KO mice. Bar graphs depict (from left to right): total distance traveled in the center zone (cm), number of entries into the center zone, total time spent in the center zone (seconds), and average velocity in the center zone (cm/sec). Data are presented as mean ± SEM for each group (WT: blue; KO: red). Statistical comparisons were performed between WT and KO groups, with significance indicated as follows: *: *p* < 0.05, **: *p* < 0.01, ***: *p* < 0.001.

### 3.9 Genotype-Specific EEG-Behavior Correlation in *Fmr1* KO Mice During Open Field Exploration

To investigate the potential relationship between EEG signal and behavioral parameters, Pearson correlations were computed between absolute and relative EEG power and OFT exploration metrics. In WT mice, no significant correlations emerged between any frequency band (absolute or relative power) and behavioral parameters **(Table IV, V, VI and VII)**. Similarly, no EEG power (absolute or relative) correlated with OFT behavior in KO mice **(Table IV, V, VI and VII)**, Except a positive correlation between absolute delta power and time spent in the center zone (r = 0.865, p = 0.008).

**Table IV.** Pearson correlation (r) and FDR-corrected p values between the absolute power of frequency band and behavioral parameters for WT mice in OFT.

|  | Distance in center |  | Entries into Center |  | Time in Center |  | Velocity |  |
| --- | --- | --- | --- | --- | --- | --- | --- | --- |
|  | r | p | r | p | r | p | r | p |
| Delta | -0.095 | 0.970 | 0.106 | 0.970 | -0.077 | 0.970 | -0.110 | 0.970 |
| Theta | 0.025 | 0.970 | 0.051 | 0.970 | -0.042 | 0.970 | 0.202 | 0.970 |
| Alpha | 0.036 | 0.970 | 0.348 | 0.970 | 0.149 | 0.970 | -0.012 | 0.970 |
| Beta | -0.097 | 0.970 | 0.114 | 0.970 | -0.088 | 0.970 | 0.144 | 0.970 |
| Gamma | -0.157 | 0.970 | -0.052 | 0.970 | -0.300 | 0.970 | 0.294 | 0.970 |
| Low Gamma | -0.203 | 0.970 | -0.109 | 0.970 | -0.308 | 0.970 | 0.291 | 0.970 |
| High Gamma | -0.065 | 0.970 | 0.058 | 0.970 | -0.271 | 0.970 | 0.286 | 0.970 |

**Table V.** Pearson correlation (r) and FDR-corrected p values between the absolute power of frequency band and behavioral parameters for *Fmr1* KO mice in OFT.

|  | Distance in center |  | Entries into Center |  | Time in Center |  | Velocity |  |
| --- | --- | --- | --- | --- | --- | --- | --- | --- |
|  | r | p | r | p | r | p | r | p |
| Delta | 0.061 | 0.916 | 0.441 | 0.575 | **0.865** | **0.008** | -0.546 | 0.575 |
| Theta | -0.147 | 0.826 | 0.251 | 0.575 | 0.538 | 0.575 | -0.460 | 0.575 |
| Alpha | -0.046 | 0.919 | 0.272 | 0.575 | 0.259 | 0.575 | -0.290 | 0.575 |
| Beta | -0.028 | 0.932 | 0.329 | 0.575 | 0.403 | 0.575 | -0.271 | 0.575 |
| Gamma | -0.073 | 0.916 | 0.317 | 0.575 | 0.406 | 0.575 | -0.289 | 0.575 |
| Low Gamma | -0.062 | 0.916 | 0.322 | 0.575 | 0.414 | 0.575 | -0.280 | 0.575 |
| High Gamma | -0.089 | 0.916 | 0.305 | 0.575 | 0.389 | 0.575 | -0.300 | 0.575 |

**Table VI.** Pearson correlation (r) and FDR-corrected p values between the relative power of frequency band and behavioral parameters for WT mice in OFT.

|  | Distance in center |  | Entries into Center |  | Time in Center |  | Velocity |  |
| --- | --- | --- | --- | --- | --- | --- | --- | --- |
|  | r | p | r | p | r | p | r | p |
| Delta | -0.107 | 0.934 | 0.021 | 0.945 | -0.117 | 0.774 | -0.172 | 0.670 |
| Theta | 0.082 | 0.934 | -0.200 | 0.766 | -0.115 | 0.774 | 0.266 | 0.670 |
| Alpha | 0.120 | 0.934 | 0.348 | 0.659 | 0.468 | 0.447 | -0.220 | 0.670 |
| Beta | -0.072 | 0.934 | -0.137 | 0.766 | -0.088 | 0.774 | 0.046 | 0.882 |
| Gamma | -0.112 | 0.934 | -0.323 | 0.659 | -0.445 | 0.447 | 0.273 | 0.670 |
| Low Gamma | -0.207 | 0.934 | -0.414 | 0.659 | -0.479 | 0.455 | 0.316 | 0.670 |
| High Gamma | 0.026 | 0.934 | -0.153 | 0.766 | 0.172 | 0.774 | 0.172 | 0.670 |

**Table VII.** Pearson correlation (r) and FDR-corrected p values between the relative power of frequency band and behavioral parameters for *Fmr1* KO mice in OFT.

|  | Distance in center |  | Entries into Center |  | Time in Center |  | Velocity |  |
| --- | --- | --- | --- | --- | --- | --- | --- | --- |
|  | r | p | r | p | r | p | r | p |
| Delta | 0.210 | 0.597 | -0.427 | 0.291 | 0.416 | 0.314 | 0.561 | 0.402 |
| Theta | -0.176 | 0.649 | -0.333 | 0.339 | -0.153 | 0.634 | 0.374 | 0.568 |
| Alpha | 0.304 | 0.469 | -0.176 | 0.585 | 0.346 | 0.378 | 0.167 | 0.503 |
| Beta | -0.036 | 0.911 | 0.505 | 0.291 | -0.255 | 0.594 | -0.007 | 0.503 |
| Gamma | -0.347 | 0.472 | 0.440 | 0.291 | -0.370 | 0.331 | -0.183 | 0.503 |
| Low Gamma | -0.308 | 0.469 | 0.443 | 0.291 | -0.354 | 0.378 | -0.162 | 0.503 |
| High Gamma | -0.374 | 0.469 | 0.389 | 0.291 | -0.200 | 0.624 | -0.200 | 0.503 |

## 4. Discussion

### 4.1 Review of EEG Phenotypes in *Fmr1* KO Models and FXS Literature

#### 4.1.1 Absolute and Relative EEG Power

This study revealed preserved absolute gamma power in juvenile female *Fmr1* KO mice on a B6 background. This pattern is distinct from the most widely reported EEG phenotype in *Fmr1* KO mice—particularly in males and on the FVB background—where increased absolute gamma power is consistently observed from juvenile through adult stages, alongside increased N1 amplitudes and sensory hypersensitivity [50–52]. For example, Lovelace et al. (2018) and Sinclair et al. (2017) reported robust increases in gamma power in both frontal and auditory cortices of male *Fmr1* KO mice, findings that closely mirror increased absolute gamma power seen in male FXS patients [23, 26–28]. On the other hand, relative gamma power was increased in the female KO mice in our study, aligning with human FXS studies that report heightened relative gamma power in both male and females [24].

Theta-band analysis revealed reduced absolute theta power in the female KO mice, though this finding shows limited consistency across preclinical and clinical studies. For instance, male *Fmr1* KO mice on the FVB background exhibited no significant genotype differences in absolute theta power during either resting-state or stimulus-evoked EEG recordings [50], while one clinical investigation noted increased absolute power across multiple frequencies (including theta) in FXS participants [22]. In contrast, relative theta power reductions observed in this study align with our recent preclinical findings of decreased relative theta power in female *Fmr1* KO mice on the FVB background, while diverging from clinical FXS data where elevated relative theta has been observed in both males and females [24, 25, 51].

Alpha oscillations exhibited the most consistent alterations across metrics in this study, with reductions in both absolute and relative alpha power. These findings mirror clinical FXS studies where diminished relative alpha power correlates with attentional deficits and cognitive impairment, with no reported sex differences in human cohorts [24, 25]. However, preclinical studies have reported divergent outcomes. For example, *Fmr1* KO rat models demonstrate reduced absolute alpha power in adult males [52], while our prior work in female *Fmr1* KO mice on the FVB background revealed increased absolute alpha power along with no change in relative alpha power [51]. Conversely, Wen et al. (2019) observed no genotype differences in absolute alpha power in male *Fmr1* KO mice while did not report relative alpha power. This variability underscores the significant influence of species, sex, and genetic background on alpha oscillatory phenotypes in FXS models.

#### 4.1.2 PAF

We observed a significant reduction in PAF in *Fmr1* KO mice during home cage recordings. While PAF has not been extensively reported in animal models, clinical studies in FXS have identified slowed alpha peaks and reduced alpha power as robust features, correlating with cognitive deficits and atypical brain maturation [22, 43]. The present findings extend this phenotype to juvenile female KO mice.

#### 4.1.3 TBR

Our results demonstrated a reduction in TBR in KO mice during home cage and light-dark test conditions. This finding contrasts with reported clinical study in FXS, which reported no significant alterations in TBR [43]. The discrepancy may reflect differences in developmental stage, sex, or species, as well as the influence of genetic background. While TBR is a well-established biomarker in ADHD and other neurodevelopmental disorders [33], its role in FXS remains less clear, and further research is needed to clarify its translational significance.

#### 4.1.4 Cross-Frequency Coupling

Analysis of cross-frequency coupling revealed reduced alpha-gamma PAC in home cage and diminished alpha-gamma and theta-gamma PAC during LD and OFT in KO mice. In clinical studies in autism spectrum disorder, altered PAC may manifest as regionally specific increases or decreases. For example, Peck et al. (2022) reported elevated resting-state alpha-gamma and theta-gamma PAC in autism participants, while other studies observed increased alpha–low gamma PAC at midline cortical sources but decreased coupling at lateral regions [54]. Though these studies focused on autism rather than FXS specifically, there could be overlapping but distinct network pathophysiology between the two.

Preclinical PAC studies in FXS models remain limited but offer critical insights. Hippocampal local field potential recordings in *Fmr1* KO mice during spatial shock avoidance tasks revealed dis-coordinated theta-gamma PAC [55]. Conversely, our prior work in female *Fmr1* KO mice on the FVB background reported elevated theta/alpha-gamma PACs, a finding opposite to the reduced PAC observed here in B6 females [5]. Thus, PAC alterations appear to be task- and region-dependent, and influenced by genetic background.

Our study also reported state-dependent alterations in AAC in juvenile female *Fmr1* KO mice, revealing a complex interplay between genotype, frequency bands, and behavioral context. During home cage recordings, KO mice exhibited increased theta–high gamma coupling alongside reduced alpha–high gamma coupling compared to WT controls. No genotype differences emerged in the LD, while OFT elicited heightened theta–low gamma AAC in KO mice. However, no consistent patterns have been observed across these conditions.

#### 4.1.5 Signal Complexity

Finally, we found no differences in EEG signal complexity, as measured by MSE, between KO and WT mice. This result differs from clinical studies reporting reduced MSE in participants (both females and males) aged 5–28 years, particularly at higher temporal scales, and is thought to reflect impaired global network integration and atypical brain maturation [43]. The preserved complexity in our juvenile female KO mice may indicate compensatory mechanisms or less severe global network disruption in this model, emphasizing the importance of considering sex and developmental stage in translational EEG research.

### 4.2 Potential Role of Thalamic GABAergic Dysfunction in Alpha Rhythm Slowing

Our study identified a selective reduction in PAF and absolute/relative alpha power in juvenile female *Fmr1* KO mice during home cage recordings. These findings align with clinical FXS studies reporting slowed alpha rhythms and reduced alpha power, which may point to a fundamental disruption in thalamocortical network function [43].

Alpha oscillations are prominent brain rhythms in the 8–13 Hz frequency range. The generation of these oscillations is a dynamic process involving both cortical and thalamic structures, with intricate thalamocortical interactions playing a central role [56, 57]. Recent evidence suggests that alpha rhythms are primarily initiated in the superficial layers of the cortex and then propagate to lower-order sensory areas, while the thalamus modulates and synchronizes these oscillations, influencing their amplitude and coherence in response to attentional and cognitive demands [57, 58]. This bidirectional communication between cortex and thalamus, mediated in part by thalamocortical relay cells, is essential for the maintenance and functional relevance of alpha activity. It is important to note, however, that while these mechanisms are characterized in humans, there are notable differences in alpha oscillation generation in mice: the dominant rhythm in the mouse is slower, and the anatomical and functional complexity of thalamocortical circuits is reduced compared to humans [59]. Therefore, caution is warranted when extrapolating findings on alpha oscillations between species.

In FXS, a range of thalamic abnormalities have been previously reported, for instance lowered fractional anisotropy, altered T-type calcium channels, and reduced thalamic gray matter density [22]. While numerous mechanisms contribute to thalamic dysfunction in FXS, two specific mechanisms have recently drawn particular attention: the dysregulation of large conductance calcium-activated potassium (BK) channels within the thalamus, and alterations in GABA_A_ receptor function in the thalamic reticular nucleus (TRN).

BK channels are widely expressed in the thalamus, where they play a complex role in shaping action potential waveforms and regulating neuronal firing frequency [62]. Recent research has demonstrated that FMRP can modulate BK channel function through two primary mechanisms: direct protein-protein interaction with the channel’s auxiliary β4 subunit, and regulation of the mRNA stability and trafficking of specific channel components [60, 61]. The direct interaction between FMRP and the β4 subunit modifies the subunit’s influence on the channel, decreasing its calcium sensitivity and thereby enhancing BK channel activity. In the absence of FMRP, as observed in FXS models, BK channel activity is diminished, leading to prolonged action potential duration, increased presynaptic calcium influx, and excessive neurotransmitter release [47, 48, 63]. Furthermore, studies have shown that inhibition of BK channels impairs thalamic burst firing dynamics, which can result in irregular alpha oscillatory activity and compromised thalamic rhythmicity [64].

GABA_A_ receptors mediate most of the fast synaptic inhibition in the TRN, generating prolonged inhibitory postsynaptic currents critical for its regulatory function [65]. Studies in both FXS patients and animal models reveal reduced GABA_A_ receptor density across multiple brain regions, including the thalamus [66–68]. FMRP interacts directly with GABA_A_ receptors to regulate their single-channel activity, modulating tonic inhibition and ensuring precise signal integration [69, 70]. Additionally, FMRP binds to mRNAs encoding GABA_A_ receptor subunits (e.g., α1, β2, γ2), stabilizing their transcripts and controlling receptor assembly [68]. In the lack of FMRP, this dual regulatory mechanism is disrupted, impairing the TRN’s ability to synchronize thalamocortical relay neurons and destabilizing thalamic rhythmicity [22]. This deficit contributes to reduced alpha oscillations, which are further exacerbated by concurrent BK channel dysfunction (prolonged burst firing) and diminished GABAergic inhibition. Together, these disruptions shift thalamocortical network activity toward lower-frequency bands, altering the balance of alpha and theta oscillations [22, 71].

Notably, while increased gamma power is a robust finding in male *Fmr1* KO mice, particularly on the FVB background, alpha rhythm slowing has not been widely reported in preclinical models, highlighting a potential sex- and strain-specific vulnerability in thalamic network function in female *Fmr1* KO mice on a B6 background. Furthermore, this mechanistic insight not only aligns our findings with clinical observations of alpha slowing and attentional deficits in FXS but also underscores the translational relevance of targeting thalamic BK channels (e.g., β4 subunit agonists) and GABA_A_ receptor pathways (e.g., benzodiazepines) as potential therapeutic strategies for restoring network synchrony in FXS.

## 5. Conclusion

This study provides a comprehensive characterization of EEG oscillatory dynamics and behavioral correlates in juvenile female *Fmr1* knockout mice on a B6 background, addressing a critical gap in the preclinical FXS literature. Our findings revealed alternation in absolute and relative EEG power, selective slowing of peak alpha frequency, and state-dependent disruptions in cross-frequency coupling. In contrast to clinical studies reporting reduced signal complexity, multiscale entropy was preserved in our model, suggesting potential compensatory mechanisms or sex- and strain-specific resilience in global network integration.

These results highlight the importance of considering sex, developmental stage, and genetic background in translational FXS research. The distinct oscillatory and behavioral phenotypes observed in this study underscore the potential need for tailored biomarker development and intervention strategies.

## List of Abbreviations Used (if any)

FXS: Fragile X Syndrome
FMR1: Fragile X Messenger Ribonucleoprotein 1
FMRP: Fragile X Messenger Ribonucleoprotein 1
B6: C57BL/6J
EEG: Electroencephalography
KO: Knockout
WT: Wildtype
PAC: Phase-amplitude coupling
AAC: Amplitude-amplitude coupling
PAF: Peak alpha frequency
TBR: Theta-beta ratio
MSE: Multi-scale entropy

## 6. Author’s contribution

BW participated in EEG surgery, recording, data analysis and interpretation, and drafted and revised the manuscript. AA participated in EEG surgery, recording, and data analysis at the beginning of the study. KM participated in data analysis and interpretation. NC designed the study, participated in data analysis and interpretation, and revised the manuscript. All authors read and approved the final manuscript.

## 7. Funding

This work was supported by the Alberta Children’s Hospital Research Foundation (NC), University of Calgary Faculty of Veterinary Medicine (NC), Natural Sciences and Engineering Research Council of Canada (NC), and FRAXA Research Foundation (NC). The funding sources had no role in the study design; in the collection, analysis, and interpretation of data; in the report’s writing; and in the decision to submit the article for publication.

## Notes

### Competing Interest Statement

The authors have declared no competing interest.

