## Supplementary Fig.1 for "Alterations in Electroencephalography Signals in Female Fragile X Syndrome Mouse Model on a C57Bl/6J Background"

### 6. Supplementary information

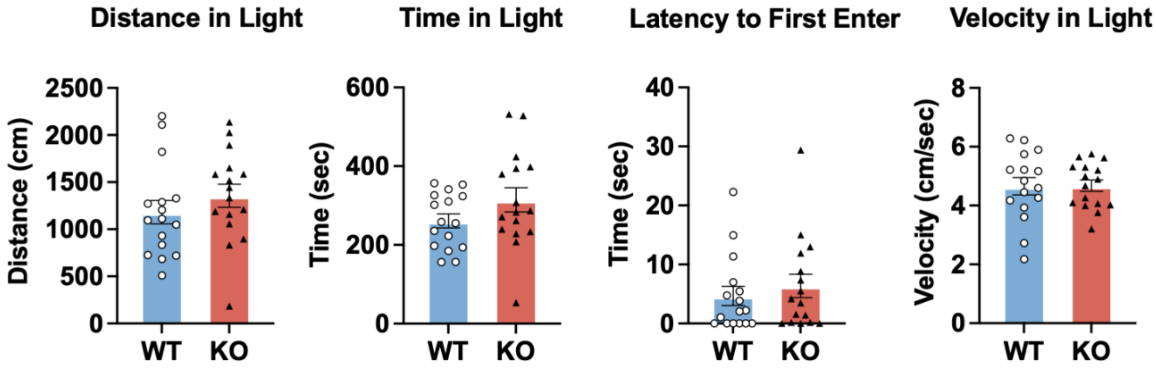


**Supplementary Figure S1. LD parameters measured in the center zone for WT and KO mice.** Bar graphs depict (from left to right): total distance traveled in the light compartment (cm), total time spent in the light compartment (seconds), latency to first enter the light compartment and average (seconds) and average velocity in the light compartment (cm/sec). Data are presented as mean ± SEM for each group (WT: blue; KO: red). Two-tailed unpaired t-tests were performed between WT and KO groups.
